# nQMaker: estimating time non-reversible amino acid substitution models

**DOI:** 10.1101/2021.10.18.464754

**Authors:** Cuong Cao Dang, Bui Quang Minh, Hanon McShea, Joanna Masel, Jennifer Eleanor James, Le Sy Vinh, Robert Lanfear

## Abstract

Amino acid substitution models are a key component in phylogenetic analyses of protein sequences. All amino acid models available to date are time-reversible, an assumption designed for computational convenience but not for biological reality. Another significant downside to time-reversible models is that they do not allow inference of rooted trees without outgroups. In this paper, we introduce a maximum likelihood approach nQMaker, an extension of the recently published QMaker method, that allows the estimation of time non-reversible amino acid substitution models and rooted phylogenetic trees from a set of protein sequence alignments. We show that the non-reversible models estimated with nQMaker are a much better fit to empirical alignments than pre-existing reversible models, across a wide range of datasets including mammals, birds, plants, fungi, and other taxa, and that the improvements in model fit scale with the size of the dataset. Notably, for the recently published plant and bird trees, these non-reversible models correctly recovered the commonly known root placements with very high statistical support without the need to use an outgroup. We provide nQMaker as an easy-to-use feature in the IQ-TREE software (http://www.iqtree.org), allowing users to estimate non-reversible models and rooted phylogenies from their own protein datasets.

## Introduction

Amino acid substitution models play an essential role in model-based phylogenetic analyses of protein sequences. Current models are typically assumed to be time reversible to ensure that model and tree estimation are computationally tractable. All time reversible models are also stationary, meaning that amino acid frequencies are at the equilibrium of the substitution matrix Q of transition rates between them. Time reversible models also obey detailed balance, i.e. fluxes between any pair of amino acids have equal magnitude in both directions. Software such as FastMG (Dang, et al., 2014) and QMaker (Minh, et al., 2021) can estimate time reversible models from collections of many multiple sequence alignments (MSAs). While mathematically convenient, there is evidence that the assumption of time reversibility may be violated (Squartini & Arndt, 2008; Naser-Khdour, et al., 2019). The challenge has been in implementing software that is computationally efficient enough to estimate time non-reversible models. If non-reversible models are a better fit to the data than reversible models, we should expect to see concomitant improvements in the estimation of tree topologies and branch lengths in phylogenetic analyses.

Another benefit of non-reversible models is that they allow the root of a phylogenetic tree to be estimated in the absence of an outgroup (Naser-Khdour, et al., 2021; Bettisworth & Stamatakis, 2021). Rooting trees is an important part of studying evolutionary relationships among species. Unfortunately, the time reversible models limit maximum likelihood (ML) methods to construct only unrooted trees since the likelihood of the tree remains the same regardless of the root position. To circumvent this limitation, most studies use outgroups to root phylogenetic trees (Maddison, et al., 1984; Huelsenbeck, et al., 2002). However, finding an appropriate outgroup for the clade under study can still a challenge in practice (Pearson, et al., 2013). Non-reversible models remove the need for an outgroup because the root position is a parameter of the model, and different rooting positions will have different likelihoods. Recent studies based on simulated and empirical data reveal encouraging results of using non-reversible models in rooting phylogenies (Naser-Khdour, et al., 2021; Bettisworth & Stamatakis, 2021).

We recently introduced QMaker (Minh, et al., 2021), a software tool that allows users to efficiently estimate reversible models from large datasets. We showed that the algorithms in QMaker improve on existing methods (Le & Gascuel, 2008; Whelan & Goldman, 2001), and used QMaker to estimate a suite of new reversible matrices that can be applied to empirical data. QMaker uses a number of approaches to make it computationally feasible to rapidly estimate new Q matrices from large collections of empirical alignments, but was restricted to estimating only time-reversible Q matrices.

In this paper, we present nQMaker, which extends QMaker to allow the estimation of stationary non-reversible models from large collections of alignments. nQMaker combines a tree search strategy to determine rooted maximum likelihood trees during the model estimation process and a ML algorithm to estimate 379 parameters of non-reversible models (instead of 179 parameters of reversible models) based on these rooted trees. We applied nQMaker to estimate six stationary non-reversible models from Pfam and five clade-specific datasets for mammals, birds, insects, yeasts, and plants. Our results show that stationary non-reversible models not only improve the fit between the model and data, but also accurately infer rooted phylogenomic trees in those cases where we had confident *a priori* knowledge of the root position from other empirical analyses.

## Material and methods

### Datasets

We used the general Pfam database (seed alignments version 31) and the same five clade-specific datasets as used in the QMaker paper (i.e., Plant, Bird, Mammal, Insect, and Yeast). The Pfam dataset consists of 13,308 MSAs from 1,150,099 sequences including 3,433,343 sites. The Pfam dataset was randomly divided into training and testing sets each containing 6,654 MSAs. The clade-specific datasets contain between 1,308 (Plant) and 7,295 (Bird) loci, and between 38 (Plant) and 343 (Yeast) sequences. For each clade-specific dataset, we randomly selected 1,000 MSAs for estimating a non-reversible model and used the remaining MSAs for testing the estimated model. We filtered out small loci with less than 50 sites in the Insect dataset (no other datasets contained loci with less than 50 sites).

The six datasets are summarized in Table 1 and available from the online supplementary material at (https://doi.org/10.6084/m9.figshare.14516712).

**Table 1.** Six datasets using for training and testing non-reversible models.

| Dataset | #Sequences | #Sites | Training | Testing | Reference |
| --- | --- | --- | --- | --- | --- |
| Pfam | 1,150,099 | 3,433,343 | 6,654 | 6,654 | (El-Gebali, et al., 2018) |
| Bird | 52 | 4,519,041 | 1,000 | 6,295 | (Jarvis, et al., 2015) |
| Insect | 144 | 595,033 | 1,000 | 1,482 | (Misof, et al., 2014) |
| Mammal | 90 | 3,050,199 | 1,000 | 3,162 | (Wu, et al., 2018) |
| Plant | 38 | 432,014 | 1,000 | 308 | (Ran, et al., 2018) |
| Yeast | 343 | 1,162,805 | 1,000 | 1,408 | (Shen, et al., 2018) |

### Methods

The amino acid substitution process is modeled by a time-homogeneous, time-continuous Markov process and represented by a 20 × 20 matrix *Q* = {*q*_*xy*_} where *q*_*xy*_ is the number of substitutions between the two different amino acids *x* and *y* per time unit (diagonal values *q*_*xx*_ are assigned such that the sum of all elements on row *x* of *Q* equals zero). In phylogenetic inference, the branch lengths reflect the number of substitutions per site, thus, the *Q* matrix is normalized by dividing the factor *μ*, where *μ* = −∑*π*_*x*_*q*_*xx*_, and *π*_*x*_ is the equilibrium frequency of 20 amino acids.

The *Q* matrix is used to calculate transition probabilities between amino acids. Specifically, the so-called transition probability matrix *P*(*t*) = {*p*_*xy*_(*t*)} where *p*_*xy*_(*t*) is the probability of changing from amino acid *x* to amino acid *y* after *t* substitutions can be calculated as follows:

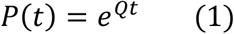

In a time-reversible model, the exchangeability rates between amino acid *x* and amino acid *y* are the same in both directions. We can only infer unrooted trees with time-reversible models because the likelihood of the tree remains the same regardless of the root placement (Felsenstein, 1981). The reversible *Q* matrix can be decomposed into a symmetric exchangeability rate matrix *R* = {*r*_*xy*_} and *Π* = {*π*_*x*_} such that *q*_*xy*_ = *π*_*y*_*r*_*xy*_ if *x* ≠ *y*, otherwise, *q*_*xx*_ = − ∑_*y*_ *q*_*xy*_. Thus, a reversible model consists of 208 free parameters (i.e., 189 parameters from the *R* matrix, and 19 parameters from *Π* vector).

If the *Q* matrix can be diagonalized, the matrix *P*(*t*) is efficiently calculated as follows:

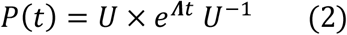

where **Λ** is the diagonal matrix of eigenvalues of *Q*; *U* is the matrix of eigenvectors of *Q* and *U*^−1^ is its inverse matrix.

In this paper, we relax the time-reversible assumption in estimating amino acid substitution models by estimating all 379 parameters of the *Q* matrix. The transition probability matrix *P*(*t*) can be calculated using a combination of eigen-decomposition and scaling-squaring techniques provided by the Eigen3 library (Guennebaud and Jacob 2010) and implemented in IQ-TREE 2 (Minh, et al., 2020). Specifically, IQ-TREE 2 uses eigen-decomposition to diagonalize *Q* into its (complex) eigenvalues, eigenvectors and inverse eigenvectors to calculate *P*(*t*) using Equation 2. If *Q* is not diagonalizable, then IQ-TREE 2 employs the scaling-squaring technique to compute *P*(*t*) based on the second order Taylor expansion of Equation 1.

Given a dataset **D** = {*D*_1_, …, *D*_*n*_} consisting of *n* multiple amino acid sequence alignments, let **T** = {*T*_1_, … *T*_*n*_} be the tree set corresponding to the dataset **D**, i.e., *T*_*i*_ is the ML tree of alignment *D*_*i*_. The ML estimation method determines the tree set **T** and a model *Q* to maximize the likelihood value *L*(*Q*, **T**; **D**). We assume that amino acid substitutions among alignments and sites are independent, thus, the likelihood value *L*(*Q*, **T**; **D**) can be calculated as follows:

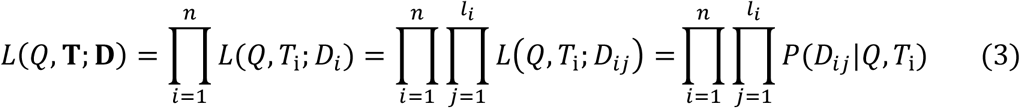

where *l*_*i*_ is the length of alignment *D*_*i*_; and *D*_*ij*_ is the data at site *j* of alignment *D*_*i*_. The likelihood value *L*(*Q, T*_i_; *D*_*ij*_) can be calculated by the conditional probability *P*(*D*_*ij*_| *Q, T*_i_) of data *D*_*ij*_ given the model *Q* and the tree *T*_*i*_.

As amino acid substitution rates vary among sites, we incorporate the site rate heterogeneity by determining site rate models ***V*** = {*V*_1_, …, *V*_*n*_} for alignments **D**, i.e., *V*_*i*_ is the site rate model of alignment *D*_*i*_. Typically, a site rate model combines a Γ distribution of rates, a proportion of invariant sites (Yang, 1993; Gu, et al., 1995), or a distribution-free rate models (Yang, 1995). The best-fit rate model for each MSA or locus was determined by using ModelFinder (Kalyaanamoorthy et al. 2017). The likelihood value *L*(*Q*, **T, V**; **D**) is now technically calculated as follows:

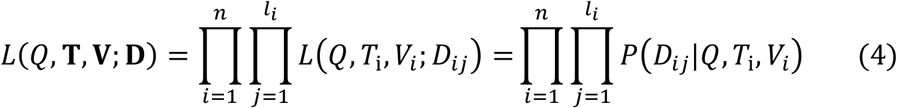

where *P*(*D*_*ij*_|*Q, T*_i_, *V*_*i*_) is the conditional probability of data *D*_*ij*_ given the model *Q*, the tree *T*_*i*_, and the site rate model *V*_*i*_.

The maximum likelihood estimation method determines parameters of the model *Q*, the trees **T** and the site rate models **V** to optimize the likelihood value *L*(*Q*, **T, V**; **D**) in Equation 4.

### Using nQMaker to estimate non-reversible models

Estimating the *Q* matrix is computationally difficult because we have to simultaneously estimate its parameters, the trees **T**, and the site rate models **V**. A number of approximate maximum-likelihood methods have been proposed to estimate model *Q* from large datasets (Minh, et al., 2021; Whelan & Goldman, 2001; Le & Gascuel, 2008; Dang, et al., 2014). The methods show that the parameters of *Q* can be accurately estimated using nearly optimal trees **T** and site rate models **V**. Thus, we can iteratively estimate the model *Q*, the trees **T**, and site rate models **V** to optimize the likelihood value *L*(*Q*, **T, V**; **D**). Currently, QMaker (Minh, et al., 2021) has been shown to efficiently estimate reversible models using this approach.

The nQMaker approach presented here extends QMaker to estimate non-reversible models from large datasets including MSAs. It composes of five main steps as illustrated in Figure 1 and described as follows:

**Figure 1:**
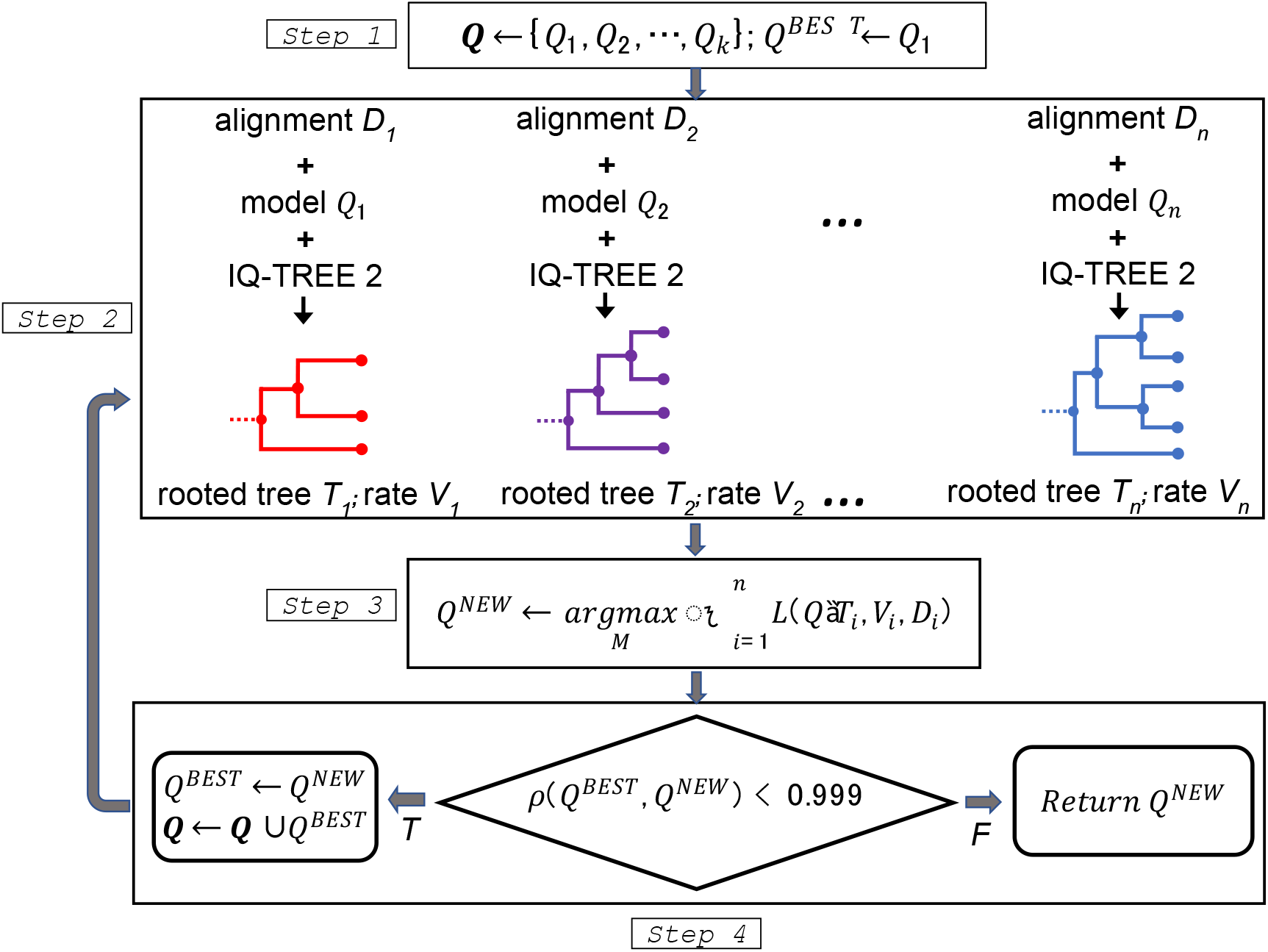
The flowchart of nQMaker to estimate a time non-reversible model from a collection of multiple protein sequence alignments.

1. Initialize a set of candidate matrices **Q**; typically we use LG (Le & Gascuel, 2008), JTT (Jones DT, 1992), and WAG (Whelan & Goldman, 2001) as three initial matrices. Set the current best matrix *Q*^*BEST*^ ≔ *LG*.
2. For each *D*_*i*_, determine *Q*_*i*_ ∈ **Q** as the best-fit matrix, *V*_*i*_ as the best site rate model, then employ IQ-TREE 2 to estimate an ML tree *T*_*i*_ based on *Q*_*i*_ and *V*_*i*_ (if *Q*_*i*_ is non-reversible, *T*_*i*_ is a rooted tree). Let 𝒯_*i*_ and ℒ_*i*_ be the topololgy and branch lengths of tree *T*_*i*_, respectively. For clade-specific datasets, instead of constructing a separate topology 𝒯_*i*_ for each locus, we estimate only one edge-linked topology 𝒯 across all loci.
3. With *V*_*i*_ and 𝒯_*i*_ fixed, estimate *Q*^*NEW*^ and ℒ_*i*_ to maximize the log-likelihood function. Precisely, we iterate two sub-steps:
  3a With *V*_*i*_, 𝒯_*i*_, and ℒ_*i*_ fixed, estimate *Q*^*NEW*^.
  3b With *V*_*i*_, 𝒯_*i*_, and *Q*^*NEW*^fixed, estimate ℒ_*i*_. If the log-likelihood is increased more than 0.1, go to step 3a, otherwise, go to the next step.
4. Assign *Q*^*BEST*^ ≔ *Q*^*NEW*^. If the Pearson correlation coefficient between *Q*^*BEST*^ and *Q*^*NEW*^ is less than 0.999, add *Q*^*BEST*^ to the set of candidate matrices **Q**, repeat from step 2. Otherwise, return *Q*^*BEST*^ as the final matrix for the database **D**.

The key difference between nQMaker and QMaker is that nQMaker uses rooted maximum likelihood trees to estimate the 379 parameters of non-reversible models, rather than using unrooted trees to estimate the 189 parameters of reversible models in QMaker. Experiments on large datasets show that the estimation process usually stops after three iterations.

### Model estimation

We used nQMaker to estimate non-reversible models (denoted NQ) from the training sets of six datasets, i.e., NQ.pfam for Pfam, NQ.plant for Plant, NQ.bird for Bird, NQ.insect for Insect, NQ.mammal for Mammal and NQ.yeast for Yeast. The reversible models for the datasets (Q.pfam, Q.plant, Q.bird, Q.insect, Q.mammal and Q.yeast) were obtained from the QMaker paper (Minh, et al., 2021). We compared non-reversible models and reversible models on testing sets using Akaike information criterion (AIC) values (Akaike, 1974). All models were tested with rate models “+G4” (Γ distribution with four categories), “+I” (invariant site model), and “+R*c*” (distribution-free rate model with *c* categories). The reversible models were also tested with “+F” option (i.e., amino acid frequencies were directly estimated from testing data). Note that each non-reversible model is represented by a single matrix *Q*, therefore “+F” option is not valid for non-reversible models.

The non-reversible model for the Pfam dataset was estimated with two commands in IQ-TREE 2:

~~~
iqtree2 -S ALN_DIR -mset LG,WAG,JTT -cmax 4
iqtree2 -S ALN_DIR.best_model.nex -te ALN_DIR.treefile --
model-joint NONREV+FO
~~~

where `-S ALN_DIR` option specifies the directory of training data; `-mset LG,WAG,JTT` option defines the initial candidate matrices to reduce computational burden; -`cmax 4` option restricts up to four categories for the rate heterogeneity across sites. The first command outputs the best models to `ALN_DIR.best_model.nex` and the best trees to `ALN_DIR.treefile`. These files are then used as the input for the second command, which estimates a join non-reversible Q matrix across all input alignments.

For clade-specific datasets, we used `-p` option instead of `-S` option to estimate an edge-linked partition model with a single tree topology shared across all loci. This `-p` option is typically used for the estimation of trees using concatenated sequences, assuming a single species tree but rescaling the branch lengths of the individual single-locus trees. Previous work has shown that edge-linked partitioned models usually perform best among among a range of related options (Duchêne, et al., 2019).

## Results

### Non-reversible models generally provided much better fit to the data than reversible models

First, we compared the non-reversible (NQ) and reversible (Q) models on the test alignments of the Pfam, bird, mammal, insect, plant and yeast datasets. Recall that the test alignments were not used to estimate the NQ matrices, ensuring that they can be used as unbiased datasets with which to compare the performance of the NQ models to other models. For each dataset, we counted the number of test alignments for which the NQ model was better than the Q model using the AIC. Table 2 shows that the NQ models fit the data better than the Q models for all clade-specific datasets, typically being selected as the best fit model for 60-70% of the test alignments. For the Pfam dataset, the reversible model Q.pfam outperformed the non-reversible model NQ.pfam, with the former being the best fit for two-thirds of the test alignments.

**Table 2.** The number of alignments where the NQ and Q models were selected as best-fit on six datasets. For example, the NQ model outperformed the Q model on 61.87% of testing alignments in the Bird dataset.

|  | **Pfam** | **Bird** | **Insect** | **Mammal** | **Plant** | **Yeast** |
| --- | --- | --- | --- | --- | --- | --- |
| *NQ* | 2218<br>(33.33%) | 3895<br>(61.87%) | 1001<br>(67.54%) | 1950<br>(61.67%) | 190<br>(61.69%) | 869<br>(61.72%) |
| *Q* | 4436<br>(66.67%) | 2400<br>(38.13%) | 481<br>(32.46%) | 1212<br>(38.33%) | 118<br>(38.31%) | 539<br>(38.28%) |

We suspected that the poor performance of NQ.pfam might be caused by a large number of small Pfam alignments (76% of Pfam test alignments have ≤ 100 sequences). This is supported by post-hoc data analysis, which shows that the NQ.pfam model outperformed the Q.pfam model in just 26% of small test alignments (with ≤ 100 sequences) but in 56% of large test alignments (with > 100 sequences). The median size of alignments best fit by NQ.pfam (78 sequences) is much larger than the median size of alignments best fit by Q.pfam (26 sequences). We further examined the effect of the number of sequences in the alignment on the model fit of NQ.pfam by classifying test alignments in Pfam into 10 subsets (bins) by the number of sequences such that *i*^*th*^ (*i* = 0 … 9) bin contains all test alignments with (*i* × 100 + 1) to (*i* × 100 + 100) sequences. We calculated the Spearman correlation between the rank of the bin and the proportion of alignments in the bin which are best fit by NQ.pfam. The Spearman correlation value is 0.903 indicating that the model fit of NQ.pfam increases with the number of sequences in testing alignments.

Second, we compared 10 different models including six non-reversible models, three general models (JTT, LG, and WAG), and one best-fit reversible model for each testing dataset (e.g. Q.pfam for Pfam or Q.plant for Plant). Similar to the results above, these results show that the non-reversible models performed best for the clade-specific datasets, but not for the Pfam dataset (Figure 2). In most cases, the second best model for each clade specific dataset was the reversible model previously estimated for that dataset (e.g. Q.mammal is the second best dataset behind NQ.mammal for the mammal dataset).

**Figure 2.**
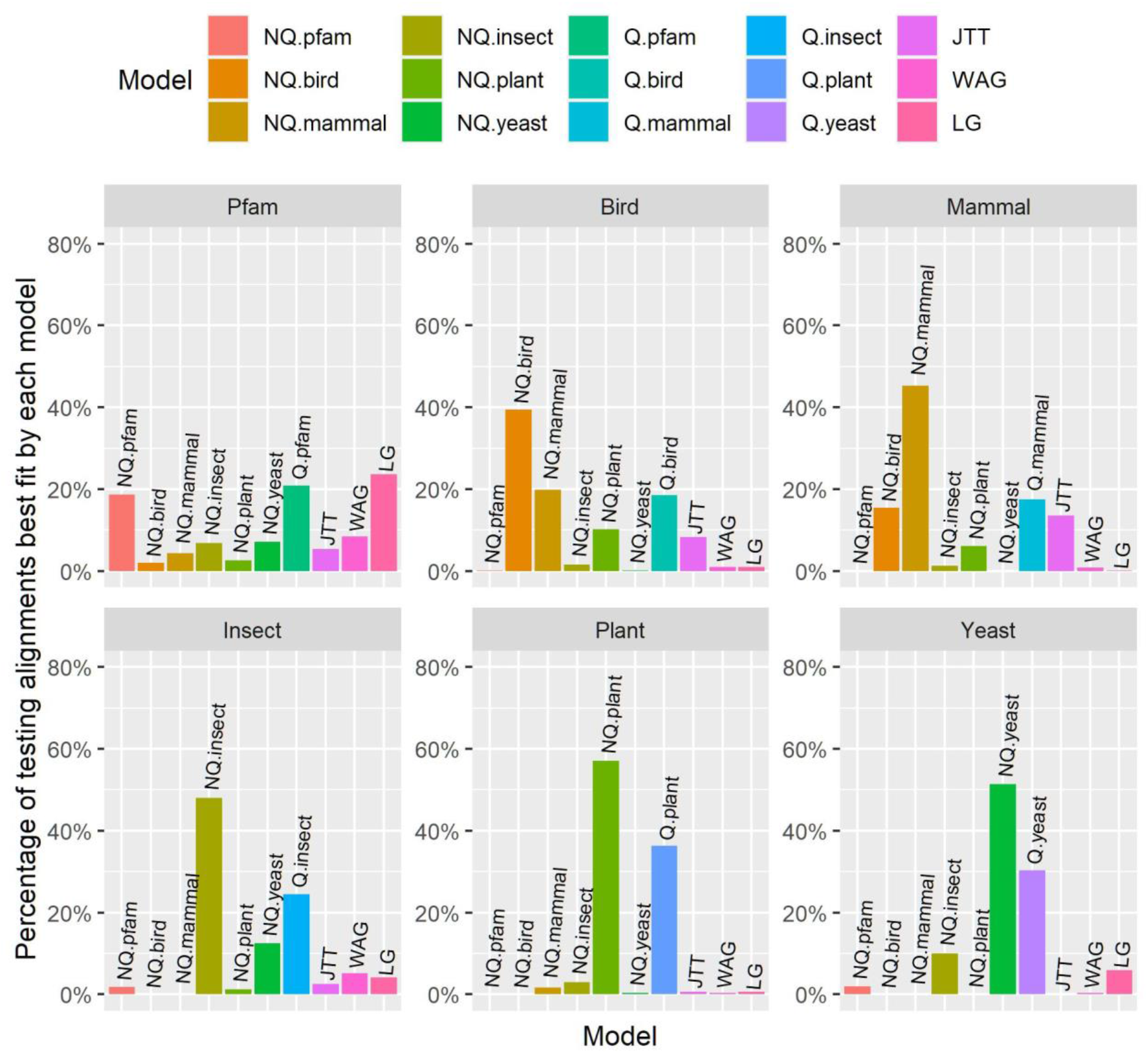
The percentage of testing alignments best fit by each model in Pfam and five clade-specific datasets.

Many genome annotations are contaminated with Pfams that do not belong to the ostensibly sequenced and assembled specie’s genome but to one of its parasites (Breitwieser, et al., 2019; Salzberg, 2019). To obtain “cleaned” clade-specific data, James et al. (James, et al., 2021) excluded all Pfam domains whose annotations suggested parasitic origin, e.g. “viral” or “transcriptase”. We used their list of cleaned (white-listed) Pfams as a filter on our training and testing Pfam sets to create a cleaned training Pfam set of 3655 MSAs and a cleaned testing Pfam set of 3611 MSAs. We chose not to use a more thoroughly cleaned version of the Pfam dataset including the removal of individual sites or sequences within each MSA as it might be too conservative and could eliminate informative data (Tan, et al., 2015). We then estimated a new non-reversible model from this cleaned Pfam dataset, which we call NQ.cPfram.

We compared the NQ.pfam model with the NQ.cPfam model estimated from the cleaned training Pfam set. Experiments showed that NQ.pfam was better than NQ.cPfam on 2519 (69.7%) out of 3611 cleaned testing MSAs. The NQ.pfam model outperformed the NQ.cPfam model on 4774 (71.7%) testing MSAs from the original Pfam dataset. Thus, the contaminated MSAs in the Pfam dataset did not considerably affect the quality of the NQ.pfam model.

### Non-reversible model fit correlates with sequence lengths

We first assessed the effect of single-locus alignment length on the model fit of NQ models on five clade-specific datasets. For each clade-specific dataset, we classified the test alignments into 10 bins by the alignment length, then calculated the Spearman correlation between the rank of the bin and the proportion of alignments which are best fit by the NQ model for that dataset. The results showed variable Spearman correlations among datasets: 0.47 for NQ.Bird, 0.87 for NQ.insect, 0.56 for NQ.Mammal, −0.02 for NQ.Plant, and 0.42 for NQ.yeast, indicating that the link between single-locus alignment length and model fit varies considerably across datasets.

We also sought to examine the fit of the new NQ models on longer concatenated alignments. To do this, we examined the model fit of NQ models on concatenated alignments from clade-specific datasets with 1, 5, 10, 20, 50, 100, and 200 loci. For each number of loci, we randomly created 100 replicate concatenated alignments, then calculated the proportion of 100 replicates where the NQ model was the best-fit model. For example, for the Plant dataset and the case of 10 loci, we created 100 concatenated alignments each composed of 10 different random loci selected from the Plant test dataset, then assessed the performance of NQ.plant on the 100 concatenated alignments. The results on five clade-specific datasets (see Figure 3) show that the proportion of replicates for which the NQ model is the best-fit model increases with the number of loci in the concatenated alignment. The NQ models outperformed the corresponding Q models on almost all concatenated alignments with ≥ 20 loci, and on practically all concatenated alignments with >50 loci (Figure 3). This result suggests that for phylogenomic datasets with many loci, non-reversible models will almost always outperform reversible models in terms of their model fit, and may therefore lead to more accurate estimation of trees and branch lengths in these cases.

**Figure 3.**
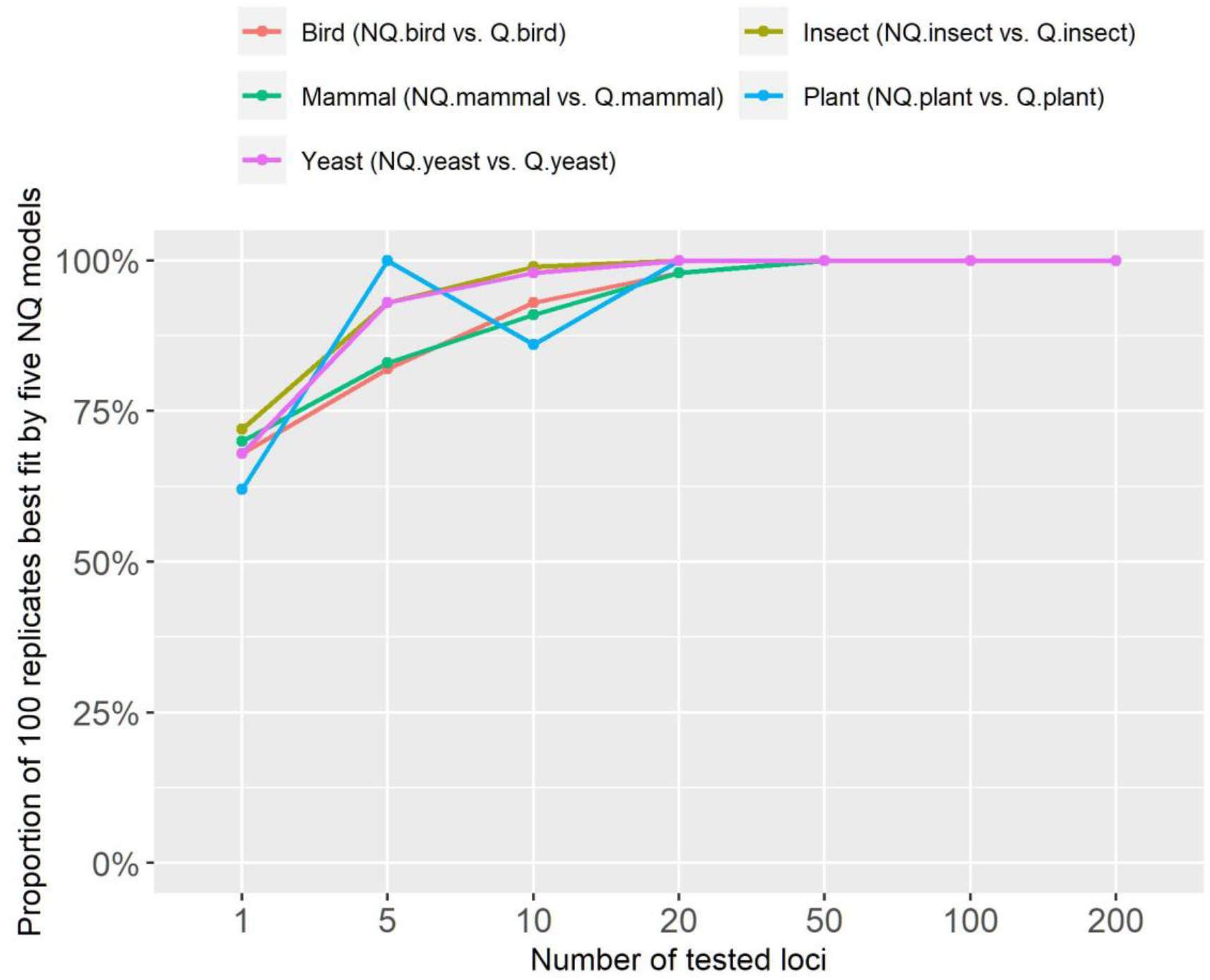
The proportion of 100 concatenated alignments best fit by non-reversible models on five clade-specific datasets.

### Analysis of the properties of non-reversible models

We used principal component analysis (PCA) to visualize the difference between non-reversible and reversible models. Each model was represented by one vector of all amino acid substitution rates and subsequently analyzed by our R script. Figure 4 illustrates the PCA analysis of six non-reversible models and 25 existing reversible models. Figure 4 shows that the models group into three distinct clusters, i.e., one cluster of non-reversible models, one cluster of reversible models estimated from mitochondrial data, and another cluster of reversible models estimate from other genomic regions. This PCA analysis indicates that non-reversible models provided a very distinct pattern of amino acid substitutions not captured by existing reversible models. To understand these NQ matrix substitution patterns, we calculated the net flux between each amino acid pair for each clade. Figure 5 shows drastic departures from reversibility in all taxonomic groups, and substantial differences between them. The largest non-reversible fluxes are not between particularly codon-adjacent or (what are typically considered) chemically-similar amino acids. Further study is needed to understand the contributions of amino acid chemistry to the direction and magnitude of the fluxes, and thus to the non-reversible evolutionary process summarized in the NQ matrices.

**Figure 4.**
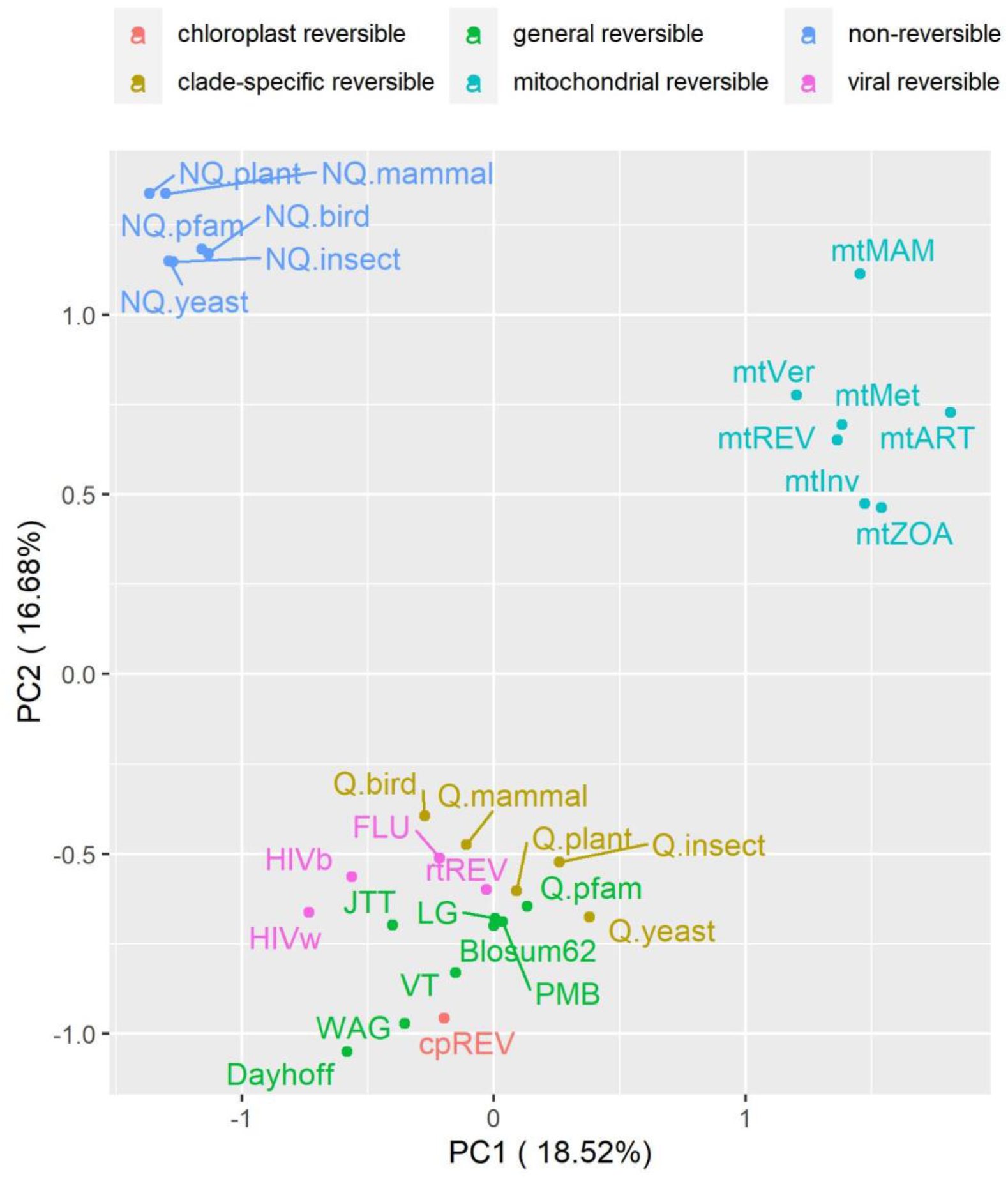
Principal component analysis of six non-reversible models and 25 reversible models. The non-reversible models are grouped into one distinct cluster.

**Figure 5.**
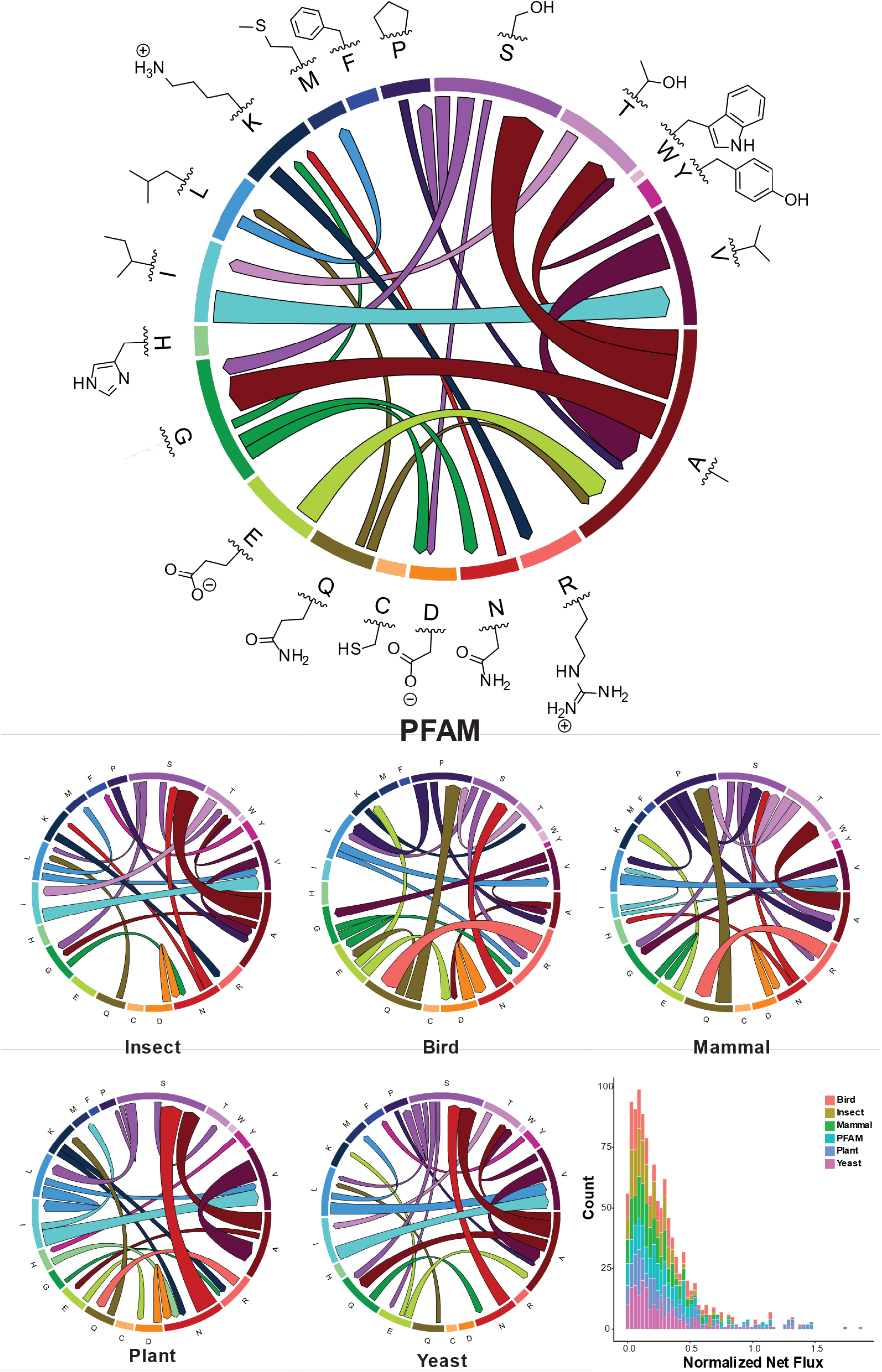
Departures from reversibility vary across taxonomic groups. Chord diagrams show net flux measurements between amino acids (represented by 1-letter codes and side-chain structures) calculated from non-reversible rate matrices, where net flux = |flux_i→j_ – flux_j→i_| = |(rate_i→j_ * freq_i._) – (rate_j→i_ * freq_j_)|. The size of each band along the outer circle represents the equilibrium frequency of each amino acid, and the width of each chord at its attachment points is proportional to the magnitude of net flux between each pair of amino acids for that taxonomic group. For clarity, only the largest 5% of net fluxes are shown. Color in chord diagrams is for ease of interpretation and contains no extra information. Inset histogram shows the distribution of all normalized net flux values for each group, each equal to (2 * net flux_ij_) / (flux_i→j_ + flux_j→i_).

### Non-reversible models correctly inferred the root placement of reconstructed trees

We assessed the root placement of trees reconstructed with non-reversible models from the two clade-specific datasets where previous publications have indicated a well-supported root placement, i.e., the plant tree from Ran et al. (Ran, et al., 2018) and the bird tree from Jarvis et al. (Jarvis, et al., 2015). The branches on reconstructed trees were labeled with rootstrap values (ranging from 0 to 1) calculated from 1000 bootstrap trees (Naser-Khdour, et al., 2021) to provide statistical support for the placement of the root on the branches. We also performed approximately unbiased (AU) test (Shimodaira, 2002) with 1000 replicates for all branches to determine a confidence set of root branches (i.e., branches with *p*_*AU*_ > 0.05 are considered as potential root branches and included into the confidence set) (Naser-Khdour, et al., 2021).

Figure 6 illustrates the plant rooted tree and the bird rooted tree reconstructed using NQ.plant and NQ.bird, respectively. The expected root branch, based on the analysis of (Ran, et al., 2018) using outgroups of the plant tree, belongs to the AU test confidence set and has a rootstrap value of 1 (supported by all bootstrap trees). Similarly, the expected root branch, based on the analysis of (Jarvis, et al., 2014) using outgroups, was confirmed by the AU test and labeled with a very high rootstrap value of 0.998 (supported by 99.8% of bootstrap trees). These results demonstrate that non-reversible models reconstructed rooted trees with high confidence in root placements that agree with the roots inferred by outgroup rooting.

**Figure 6.**
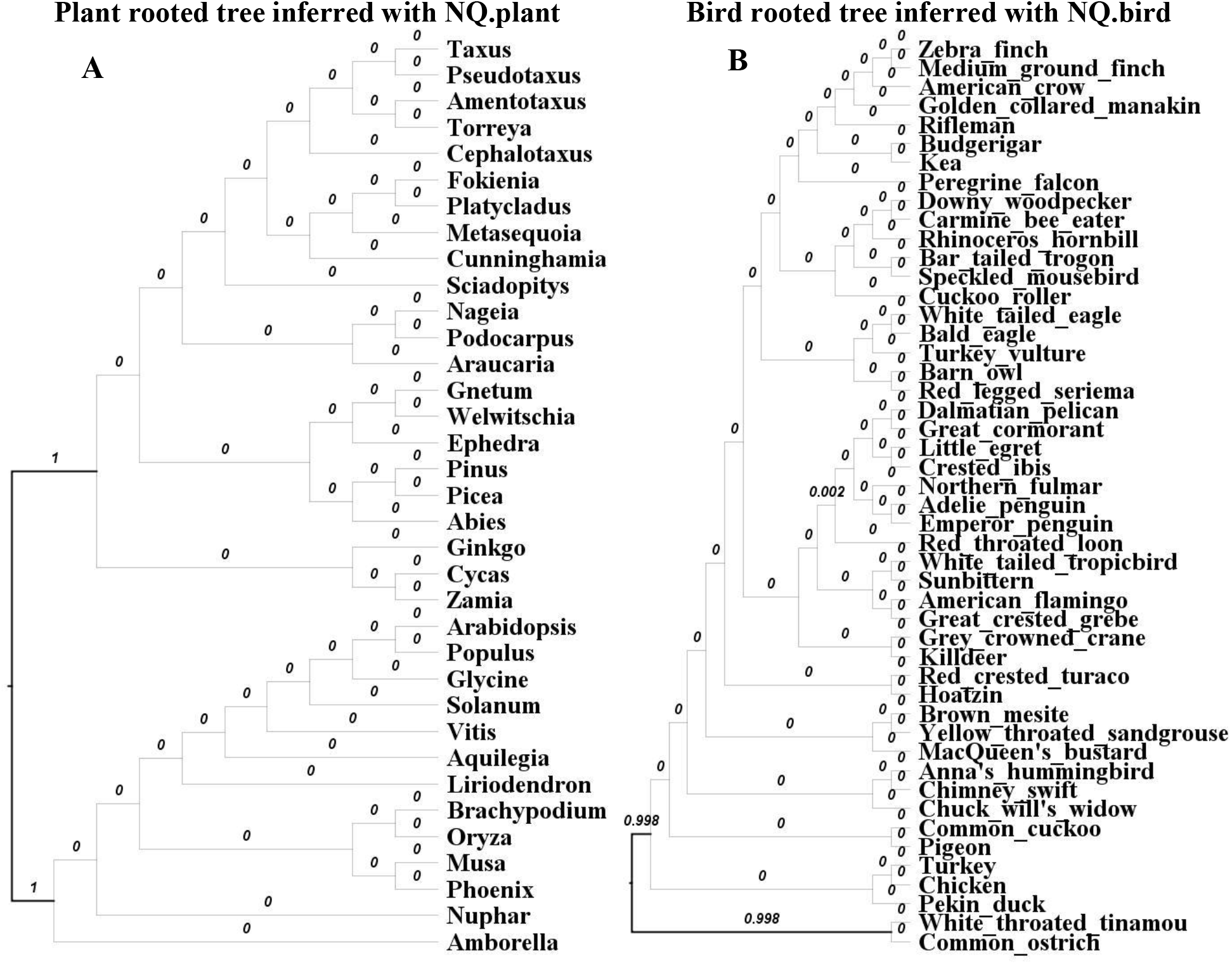
The plant rooted tree of 35 species (A) reconstructed from a concatenated protein alignment of 1308 loci using IQ-TREE 2 with the NQ.plant model. The bird rooted tree of 48 species (B) reconstructed from a concatenated protein alignment of 8295 loci using the NQ.bird model. Bold branches are branches contained in the confidence set of the AU test and numbers displaying on branches are the rootstrap values.

### Non-reversible models inferred different locus trees and coalescent based species trees

Next, we examined whether the six new non-reversible matrices can infer different tree topologies. For each single-locus MSA in each dataset, we inferred an unrooted ML tree using the best-fit model among nine published reversible models (JTT, WAG, LG, Q.pfam, Q.plam, Q.mammal, Q.bird, Q.insect and Q.yeast), which we call T_REV_. We then performed a second IQ-TREE run considering 15 models, comprising the same nine reversible models but adding the six new non-reversible models (NQ.pfam, NQ.plant, NQ.mammal, NQ.bird, NQ.insect, or NQ.yeast), to infer another tree T_NEW_. If one of the six NQ models fits the data better, then T_NEW_ will be rooted and will therefore differ from T_REV_. In this case we launch another IQ-TREE run with a same matrix as T_REV_ but using a different random seed. We call the resulting tree T_REV2_. Otherwise, if NQ models do not provide a better fit, then the 2^nd^ run will use the same model as the first run but T_NEW_ might still be different from T_REV_ due to search heuristics. Thus, for each alignment we now have three trees T_REV_, T_NEW_, and T_REV2_ when a non-reversible model fits the data best.

We then compared the three trees for each alignment when a non-reversible model fits the data best using normalized Robinson-Foulds (nRF) distances. To calculate the nRF we first unrooted the rooted tree (if required) then used IQ-TREE to calculate the nRF with options -rf1 --normalize-dist. To ask whether non-reversible models lead to bigger changes in tree topologies than expected from search heuristics alone, we compared the two distributions of normalized Robinson-Foulds (nRF) (Robinson and Foulds 1981) nRF(T_NEW_, T_REV_) and nRF(T_REV_, T_REV2_). The two nRF distributions are depicted in Figure 7. We found that using non-reversible models changes locus tree topologies in every dataset (the red line) and, particularly in the Pfam dataset, changes are somewhat greater between reversible and non-reversible models than between reversible models initiated with different random seeds (the blue line).

**Figure 7.**
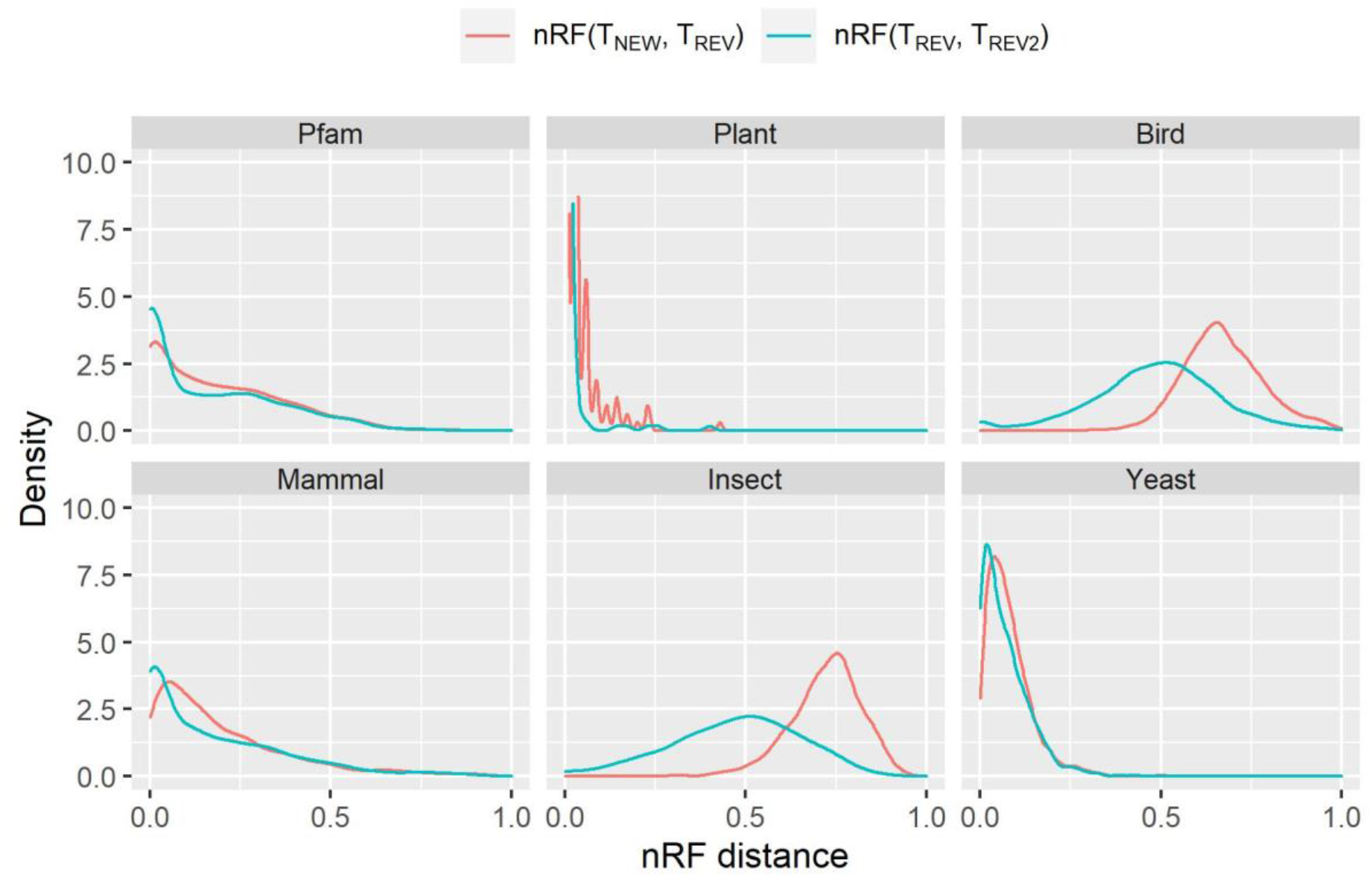
Distributions of normalized Robinson-Foulds (nRF) distances between the trees inferred by non-reversible and reversible models. The red line is the distribution where the best-fit model is one of the new non-reversible models inferred in this study (NQ.pfam, NQ.plant, NQ.mammal, NQ.bird, NQ.insect, or NQ.yeast). Comparing to best-fit reversible model, new model shows an effect on the tree topology (the best-fit reversible model is chosen from nine existing models Q.pfam, Q.plant, Q.mammal, Q.bird, Q.insect, Q.yeast, LG, JTT, or WAG; and is showed by the blue line).

Because of the observed differences between gene tree topologies, we examined to what extent it influences the reconstruction of species trees using coalescent based methods. These methods use distributions of single-locus trees to infer a species tree, so changes in the underlying single-locus trees may affect species-tree inference. To this end, for each clade-specific dataset, we used ASTRAL version 5.15 (Zhang, et al., 2018) to construct a species tree ASTRAL_REV_ from the set of T_REV_ and a species tree ASTRAL_NEW_ from the set of T_NEW_ trees. For plant dataset, the ASTRAL_REV_ tree and the ASTRAL_NEW_ tree (Figure 8A) differ by the position of a single taxon, Liriodendron. The topological differences are more pronounced for Mammals, Insects, Yeasts, Birds with 2, 10, 15, and 17 different branches between the ASTRAL_REV_ and ASTRAL_NEW_ trees. Figure 8B highlights these differences for the Bird dataset.

**Figure 8.**
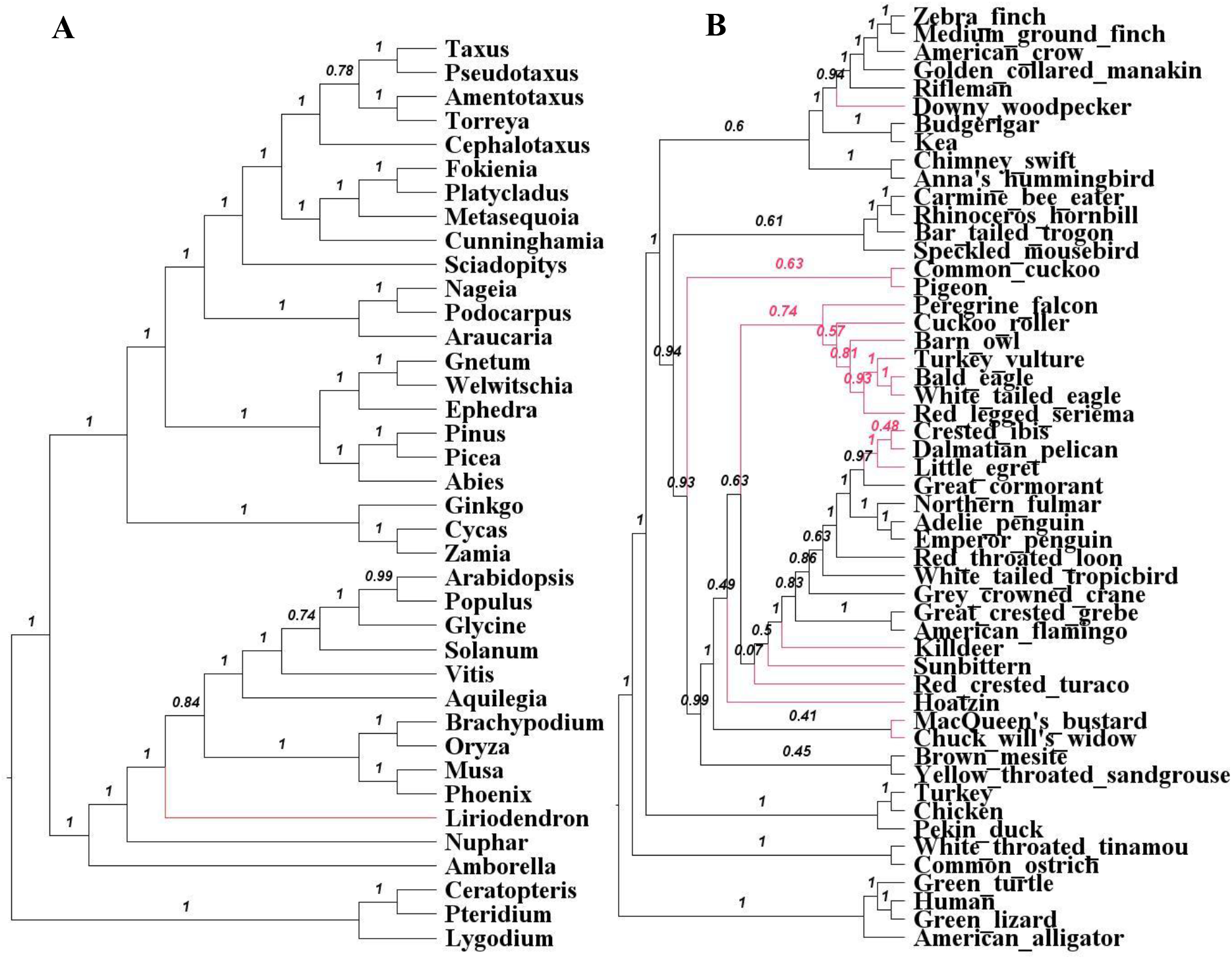
ASTRAL_NEW_ species trees from Plant (A) and Bird (B) data reconstructed from the set of T_NEW_ locus trees. Shown on each internal branch the ASTRAL local posterior probability.

## Discussion

Most phylogenetic analyses of protein sequences use time-reversible substitution models, which can be limited in their ability to accurately model the biological process of amino acid substitution. Although estimating time non-reversible models is complicated and computationally expensive (e.g., 105 days with a computer of 36 cores for estimating NQ.pfam), it has the potential to allow model of sequence evolution to better reflect the underlying evolutionary mechanisms, and hence could improve the estimation of evolutionary relationships and timescales among species.

In this paper, we introduced a new approach, nQMaker, to estimate non-reversible models from large datasets including hundreds to thousands of MSAs. We applied nQMaker to estimate six non-reversible models: a general protein model from Pfam and five clade-specific datasets for birds, insects, mammals, plants, and yeasts respectively. Our analyses show that the non-reversible models capture a distinct pattern of amino acid substitutions not captured by the traditional reversible models, that the non-reversible models affect the inference of tree topologies, and allow for the estimation of root positions without outgroups.

Our results show that non-reversible models are often selected in preference to reversible models, and that this tendency increases with the size of the alignment. Non-reversible models were selected using standard model selection approaches for most single-locus alignments. In concatenated multi-locus alignments, non-reversible models tended to be the best fit model in practically all datasets with at least 20 loci. The trees inferred with non-reversible models were often topologically different from those constructed with reversible models, suggesting that when a non-reversible model is the best-fit model for a dataset, topological accuracy of phylogenetic inference may be improved.

Rooting phylogenetic trees is an essential task in studying evolutionary relationships among species. This is normally accomplished by using outgroup species or additional assumptions such as molecular clocks (Huelsenbeck, et al., 2002). Non-reversible models provide an alternative approach that implicitly enables the reconstruction of rooted trees as part of the model. Our analyses of Bird and Plant datasets with non-reversible models identified the root of the trees of these groups with a very high statistical confidence that agree with previous studies (Ran, et al., 2018; Jarvis, et al., 2015). Together with other encouraging results on mammals (Naser-Khdour, et al., 2021) and from simulated data (Bettisworth & Stamatakis, 2021), this provides increasing evidence that non-reversible models are effective and accurate in identifying root placements for empirical datasets, and will especially be useful when an appropriate outgroup is difficult to obtain.

The non-reversible models consist of 379 parameters, the pairwise substitution rates between 20 amino-acids. Therefore, they should be estimated from large datasets consisting of hundreds to thousands MSAs to avoid over-fitting the data. The six non-reversible rate matrices we estimate in this study are now available in the latest version of IQ-TREE 2, allowing researchers to readily utilize these models for their datasets. We recommend that users perform model selection to determine the best fit model for any specific alignment under study, and note that it is possible to combine both reversible and non-reversible models in a single partitioned analysis. The nQMaker algorithm is implemented in IQ-TREE 2, so researchers can estimate non-reversible models from their own datasets. For example, the NQ.plant model was estimated from 1000 plant alignments in 1.5 days using a computer with 36 cores.

A limitation of our models is that while relaxing the time reversibility, they still assume stationarity, i.e., the amino acid frequencies stay constant along the tree. However, the stationary assumption is highly likely to be violated during the evolution of distantly related proteins, e.g., between bacteria and eukaryotes. Failure to account to heterogeneous sequence composition might mislead phylogenetic reconstruction. Apart from non-stationary models, one can also use a mixture model of several Q matrices such as C10-C60, LG4M and LG4X (Le, et al., 2012). Therefore, deriving non-stationary and/or mixture amino acid models will be an important avenue of future research.

## References

Akaike, H., 1974. A new look at the statistical model identification. IEEE Trans Autom Control, p. 19:716–23.

Bettisworth, B. & Stamatakis, A., 2021. Root Digger: a root placement program for phylogenetic trees. BMC Bioinformatics, Volume 22, p. 225.

Breitwieser, F. P., Pertea, M., Zimin, A. V. & Salzberg, S. L., 2019. Human contamination in bacterial genomes has created thousands of spurious proteins.. Genome research, 6, 29(6), pp. 954–960.

Dang, C. C. et al., 2014. FastMG: a simple, fast, and accurate maximum likelihood procedure to estimate amino acid replacement rate matrices from large data sets. BMC Bioinformatics, Volume 15, p. 341.

Duchêne, D. A. et al., 2019. Linking Branch Lengths across Sets of Loci Provides the Highest Statistical Support for Phylogenetic Inference. Molecular Biology and Evolution, 12, Volume 37, pp. 1202–1210.

El-Gebali, S. et al., 2018. The Pfam protein families database in 2019. Nucleic Acids Research, 10, Volume 47, pp. D427–D432.

Felsenstein, J., 1981. Evolutionary trees from DNA sequences: A maximum likelihood approach. Journal of Molecular Evolution, Volume 17, p. 368–376.

Gu, X., Fu, Y.-X. & Li, W.-H., 1995. Maximum likelihood estimation of the heterogeneity of substitution rate among nucleotide sites. Molecular Biology and Evolution, 12(4), p. 546–557.

Huelsenbeck, J. P., Bollback, J. P. & Levine, A. M., 2002. Inferring the Root of a Phylogenetic Tree. Systematic Biology, 1, Volume 51, pp. 32–43.

James, J. E. et al., 2021. Universal and taxon-specific trends in protein sequences as a function of age.. eLife, 1.Volume 10.

Jarvis, E. D. et al., 2014. Whole-genome analyses resolve early branches in the tree of life of modern birds. Science, Volume 346, p. 1320–1331.

Jarvis, E. D. et al., 2015. Phylogenomic analyses data of the avian phylogenomics project. GigaScience, 2.Volume 4.

Jones DT, T. W. T. J., 1992. The rapid generation of mutation data matrices from protein sequences. Bioinformatics, 8(3), pp. 275–282.

Le, S. Q., Dang, C. C. & Gascuel, O., 2012. Modeling Protein Evolution with Several Amino Acid Replacement Matrices Depending on Site Rates. Molecular Biology and Evolution, 4, Volume 29, pp. 2921–2936.

Le, S. Q. & Gascuel, O., 2008. An improved general amino acid replacement matrix. Molecular Biology and Evolution, p. 25:1307–20.

Maddison, W. P., Donoghue, M. J. & Maddison, D. R., 1984. Outgroup Analysis and Parsimony. Systematic Biology, 3, Volume 33, pp. 83–103.

Minh, B. Q., Dang, C. C., Vinh, L. S. & Lanfear, R., 2021. QMaker: Fast and accurate method to estimate empirical models of protein evolution. Systematic Biology.

Minh, B. Q. et al., 2020. IQ-TREE 2: New models and efficient methods for phylogenetic inference in the genomic era. Molecular Biology and Evolution, 11, 37(5), p. 1530–1534.

Misof, B. et al., 2014. Phylogenomics resolves the timing and pattern of insect evolution. Science, Volume 346, p. 763–767.

Naser-Khdour, S., Minh, B. Q. & Lanfear, R., 2021. Assessing Confidence in Root Placement on Phylogenies: An Empirical Study Using Non-Reversible Models for Mammals. Systematic Biology.

Naser-Khdour, S. et al., 2019. The Prevalence and Impact of Model Violations in Phylogenetic Analysis. Genome Biology and Evolution, 9, Volume 11, pp. 3341–3352.

Nguyen, L.-T., Schmidt, H. A., von Haeseler, A. & Minh, B. Q., 2014. IQ-TREE: A Fast and Effective Stochastic Algorithm for Estimating Maximum-Likelihood Phylogenies. Molecular Biology and Evolution, 11, Volume 32, pp. 268–274.

Pearson, T. et al., 2013. When Outgroups Fail; Phylogenomics of Rooting the Emerging Pathogen, Coxiella burnetii. Systematic Biology, 7, Volume 62, pp. 752–762.

Ran, J.-H., Shen, T.-T., Wang, M.-M. & Wang, X.-Q., 2018. Phylogenomics resolves the deep phylogeny of seed plants and indicates partial convergent or homoplastic evolution between Gnetales and angiosperms. Proceedings of the Royal Society B: Biological Sciences, Volume 285, p. 20181012.

Robinson, D. F. & Foulds, L. R., 1981. Comparison of phylogenetic trees. Mathematical Biosciences, Volume 53, pp. 131–147.

Salzberg, S. L., 2019. Next-generation genome annotation: we still struggle to get it right. Genome Biology, Volume 20, p. 92.

Sayyari, E., Whitfield, J. B. & Mirarab, S., 2017. Fragmentary Gene Sequences Negatively Impact Gene Tree and Species Tree Reconstruction. Molecular Biology and Evolution, 10, Volume 34, pp. 3279–3291.

Shen, X.-X. et al., 2018. Tempo and Mode of Genome Evolution in the Budding Yeast Subphylum. Cell, Volume 175, pp. 1533–1545.e20.

Shimodaira, H., 2002. An Approximately Unbiased Test of Phylogenetic Tree Selection. Systematic Biology, 5, Volume 51, pp. 492–508.

Squartini, F. & Arndt, P. F., 2008. Quantifying the Stationarity and Time Reversibility of the Nucleotide Substitution Process. Molecular Biology and Evolution, 8, Volume 25, pp. 2525–2535.

Tan, G. et al., 2015. Current Methods for Automated Filtering of Multiple Sequence Alignments Frequently Worsen Single-Gene Phylogenetic Inference. Systematic Biology, 6, Volume 64, pp. 778–791.

Whelan, S. & Goldman, N., 2001. A General Empirical Model of Protein Evolution Derived from Multiple Protein Families Using a Maximum-Likelihood Approach. Molecular Biology and Evolution, 5, Volume 18, pp. 691–699.

Wu, S., Edwards, S. & Liu, L., 2018. Genome-scale DNA sequence data and the evolutionary history of placental mammals. Data in Brief, Volume 18, pp. 1972–1975.

Yang, Z., 1993. Maximum-likelihood estimation of phylogeny from DNA sequences when substitution rates differ over sites. Molecular Biology and Evolution, pp. 10:1396-1401.

Yang, Z., 1995. A space-time process model for the evolution of DNA sequences. Genetics, Volume 139, p. 993–1005.

Zhang, C., Rabiee, M., Sayyari, E. & Mirarab, S., 2018. ASTRAL-III: polynomial time species tree reconstruction from partially resolved gene trees. BMC Bioinformatics, Volume 19, p. 153.

